# Warmer world gets sicker – meta-analysis reveals strong increase in parasitism at elevated temperatures across diverse host-parasite systems

**DOI:** 10.64898/2026.03.16.712045

**Authors:** Adam Z. Hasik, Slawek Cerbin, Justyna Wolinska

## Abstract

Climate change is expected to alter infectious disease dynamics worldwide, yet evidence that rising temperatures increase infection (warmer, sicker world hypothesis; WSWH) remains equivocal. We conducted a meta-analysis of 775 effect sizes from 124 experimental studies spanning terrestrial, freshwater, and marine host−parasite systems to test whether warming increases parasitism. Elevated temperatures increased infection, producing moderate positive effects on prevalence and infection intensity in non-phylogenetic models. Temperature effects were significant in terrestrial systems and varied markedly among taxa. Strong responses were observed in nematodes and fungal parasites as well as in plant, invertebrate, and bacterial hosts, whereas vertebrates showed little evidence of consistent increases. Phylogenetically-controlled analyses yielded similar but less precise estimates, indicating that responses are unevenly distributed across the Tree of Life. Our synthesis provides the most comprehensive experimental test of the WSWH to date and shows that warming can elevate infection risk while generating heterogeneous outcomes among host−parasite systems.

## Introduction

The negative impacts of anthropogenic climate change on the natural world have been the focus of ecological and evolutionary studies for decades^1,2^. Rising temperatures and increasing climatic variability threaten species that are unable to physiologically or behaviorally cope with these changes, often resulting in range shifts, global extirpations, or extinctions^3,4^. While the direct effects of climate change on free-living species are well documented, a less studied aspect is how rising temperatures may indirectly affect these organisms through parasite-mediated interactions.

Given the multifaceted nature of climate change^5-7^ and the system-specific responses of host-parasite interactions^8^, it can be difficult to directly link increasing temperatures to elevated parasitism. Nevertheless, accumulating evidence across taxa suggests that rising temperatures can increase parasitism, an idea formalized in the *warmer, sicker world hypothesis* (WSWH)^9^. The WSWH posits that increased temperatures will, on average, increase infection rates and disease risk. The WSWH is based on the fact that many parasites are small-bodied organisms with developmental rates and life-cycles that often accelerate with warming, potentially increasing transmission, infection rates, and disease burden at broad spatial scales.

It has been over 30 years since the WSWH was first proposed^9-13^, yet evidence for its generality remains mixed despite several syntheses and reviews. Meta-analyses have offered contrasting conclusions: one study of 378 effect sizes reported a moderate positive association between climate change and infection, with mean temperature as the only significant predictor and no detectable effects of precipitation or climate variability^14^; another, analyzing 131 interactions examining terrestrial animal hosts, detected no consistent temperature effects^15^. Similarly, tests of the thermal mismatch hypothesis, which predicts elevated parasitism when hosts experience conditions outside of their thermal optimum, found warming increased parasitism only in cool-adapted hosts^16,17^, challenging the universal predictions of the WSWH.

While these studies represent important advances, several limitations remain. Previous syntheses have largely focused on terrestrial^15,16^ or freshwater systems^16^ and often did not control for phylogenetic relationships among hosts and parasites^14,16^. Further, the vast majority of evidence evaluating the WSWH derives from observational studies^14-16^, which limits inference about causality. Given the nuanced and context-dependent nature of temperature effects on parasitism, more detailed examination from experimental studies is needed, where explicit links between the effect of increased temperatures and rates of parasitism can be made.

Here, we present a comprehensive, phylogenetically-controlled meta-analysis of experimental studies spanning terrestrial, freshwater, and marine systems. By focusing exclusively on temperature manipulations, we provide a rigorous test of the WSWH. Specifically, we assessed whether warming directly increases parasitism, rather than relying on correlative patterns from observational studies. We synthetized data comparing parasitism metrics - prevalence (proportion of hosts infected) and intensity (mean number of parasites per infected host) - between control and elevated-temperature treatments. To account for evolutionary non-independence, we incorporated phylogenetic relationships among both hosts and parasites using the Open Tree of Life^18^, and included host-parasite phylogenetic interactions^8,15,19^. Because not all taxa could be mapped phylogenetically, we constructed two datasets: a phylogenetic dataset (654 effect sizes from 102 studies) and an overall dataset including unmapped taxa and viral parasites (775 effect sizes from 124 studies). We conducted three complementary analyses: (i) controlling for phylogeny, reduced phylogenetic dataset; (ii) not controlling for phylogeny, reduced phylogenetic dataset; and (iii) not controlling for phylogeny, full dataset. Comparisons between analyses (i) and (ii) assessed the influence of shared evolutionary history, while comparisons between (ii) and (iii) evaluated the inclusion of viral parasites. Effects of warming were quantified using Hedge’s *g* (standardized mean difference), with positive values indicating increased parasitism. Hedge’s *g* values of 0 – 0.3, 0.3 – 0.7, and > 0.7 correspond to small, moderate, and large effects, respectively^20^.

Our meta-analysis revealed multiple positive effects of warming on parasitism, robust to both phylogenetic correction and viral inclusion. Beyond experimentally supporting the WSWH, our findings highlight taxonomic and geographic gaps in the literature and underscore the need for further experimental work in understudied host-parasite systems.

## Results

### General patterns

Across all model-dataset combinations, warming generally increased parasitism. In the phylogenetically-controlled model excluding viral parasites (analysis i), the estimated effect was large but not statistically-significant (effect size [95% CI]: *θ* = 0.89 [-0.29, 2.06]; Fig. 1a). Models that did not account for phylogeny (analyses ii and iii) revealed moderate, statistically-significant positive effects, both when viruses were excluded (*θ* = 0.53 [0.24, 0.83]; Fig. 1b) and included (*θ* = 0.46 [0.21, 0.71]; Fig. 1c).

**Fig. 1.**
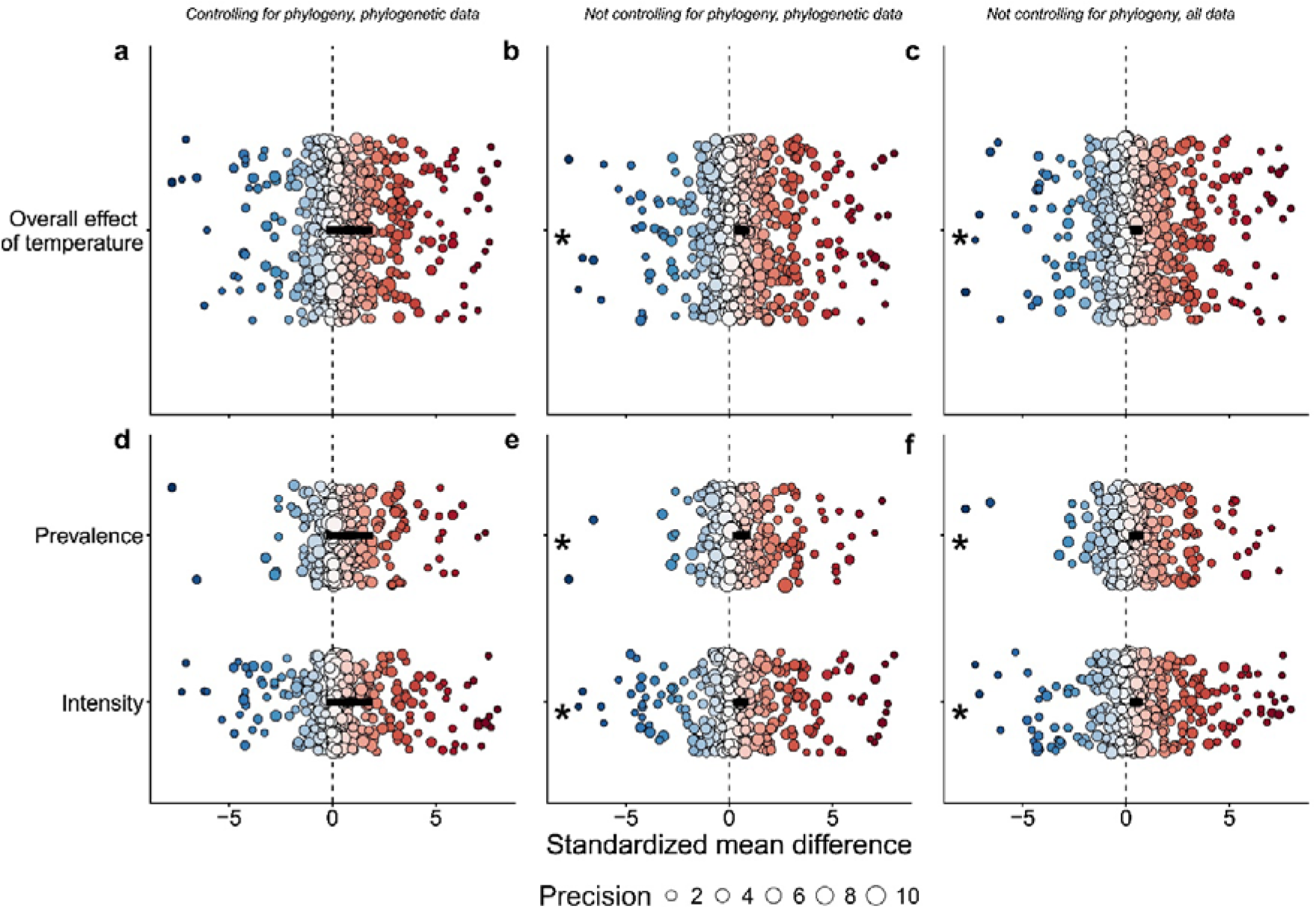
Overall effects of temperature on parasitism. Effects of elevated temperature across model structures. Columns = model type (phylogenetic / non-phylogenetic same dataset / non-phylogenetic full dataset); rows = parasitism measure (overall/top, prevalence & intensity/bottom). Effect sizes are truncated to −8 to 8 (full range: −29.59 to 102.12). Positive values indicate increased parasitism under warming (negative = reduced). Black squares show model estimates (±95% CI). Points show individual effect sizes, colored by magnitude and sign (cooler = more negative; warmer = more positive) and scaled by precision (1/SE). Asterisks denote statistically-significant estimates.

Separating parasitism into prevalence and intensity yielded consistent patterns. In the phylogenetically-controlled model, effect sizes were larger but not statistically-significant (*θ*_*prevalence*_ = 0.91 [-0.29, 2.11]; *θ*_*intensity*_ = 0.87 [-0.31, 2.06]; Fig. 1d). Non-phylogenetic models revealed significant warming-driven increases in both metrics, regardless of viral inclusion (viruses excluded: *θ*_*prevalence*_ = 0.58 [0.19, 0.97]; *θ*_*intensity*_ = 0.51 [0.17, 0.84]; Fig. 1e; viruses included: *θ*_*prevalence*_ = 0.47 [0.15, 0.80]; *θ*_*intensity*_ = 0.45 [0.16, 0.73]; Fig. 1f).

### Effects of parasite reproductive strategy and taxonomic group

No significant effects were detected in the phylogenetically-controlled analysis (Fig. 2a, Table S1) whereas in non-phylogenetic models, both microparasites and macroparasites exhibited significant increases in parasitism under warming (Fig. 2b-c). Analyses by parasite taxonomic group revealed substantial heterogeneity. Nematodes were particularly temperature-sensitive, with consistently large effect sizes (all *θ* > 2) across all models (Fig. 2d-f). Fungal parasites also exhibited positive temperature effects in the non-phylogenetic full dataset analysis (*θ* = 0.52 [0.02, 1.01], Fig. 2f). Other parasite groups showed variable, non-significant responses (Fig. 2d-f; Table S1).

**Fig. 2.**
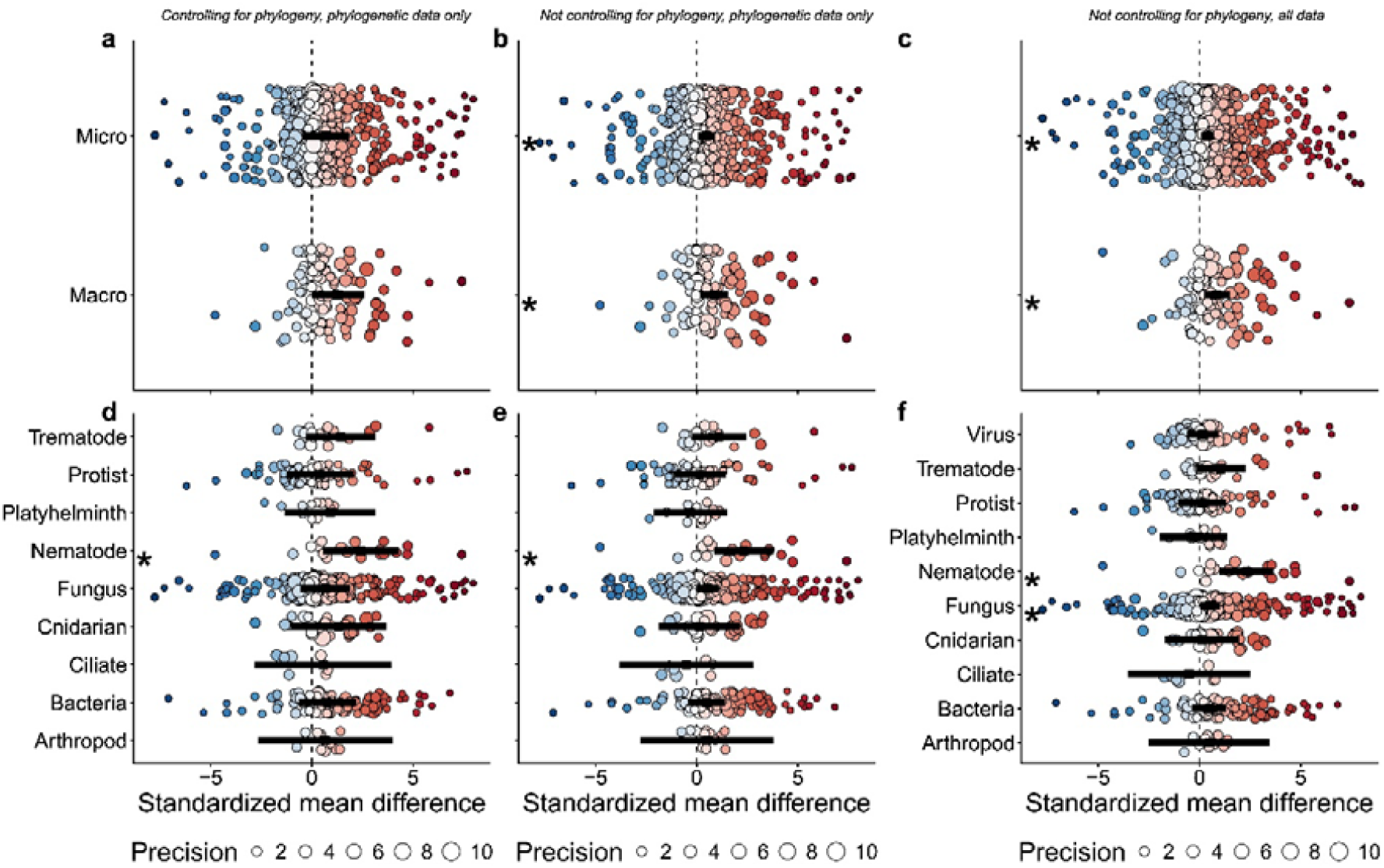
Temperature effects by parasite reproductive strategy and taxonomic group. Effects of elevated temperature partitioned by parasite reproductive strategy (top: microparasites, macroparasites) and taxonomic group (bottom). Columns = model type (phylogenetic, non-phylogenetic, full dataset). Effect sizes are truncated to −8 to 8 (full range: −29.59 to 102.12). Positive values indicate increased parasitism under warming. Black squares show model estimates (±95% CI). Points show individual effect sizes, colored by magnitude and scaled by precision (1/SE). Asterisks denote statistically-significant estimates.

### Effects of host type

Host responses differed markedly across taxa. Plants, invertebrates, and bacterial hosts experienced significant increases in parasitism with warming, with effect sizes consistently exceeding *θ* > 0.65 for plants and invertebrates and *θ* > 2.7 for bacteria across all analyses (Fig. 3a-c). In contrast, vertebrate hosts showed no significant temperature-parasitism relationship, with estimated mean effects slightly negative (Fig. 3a-c, Table S2).

**Fig. 3.**
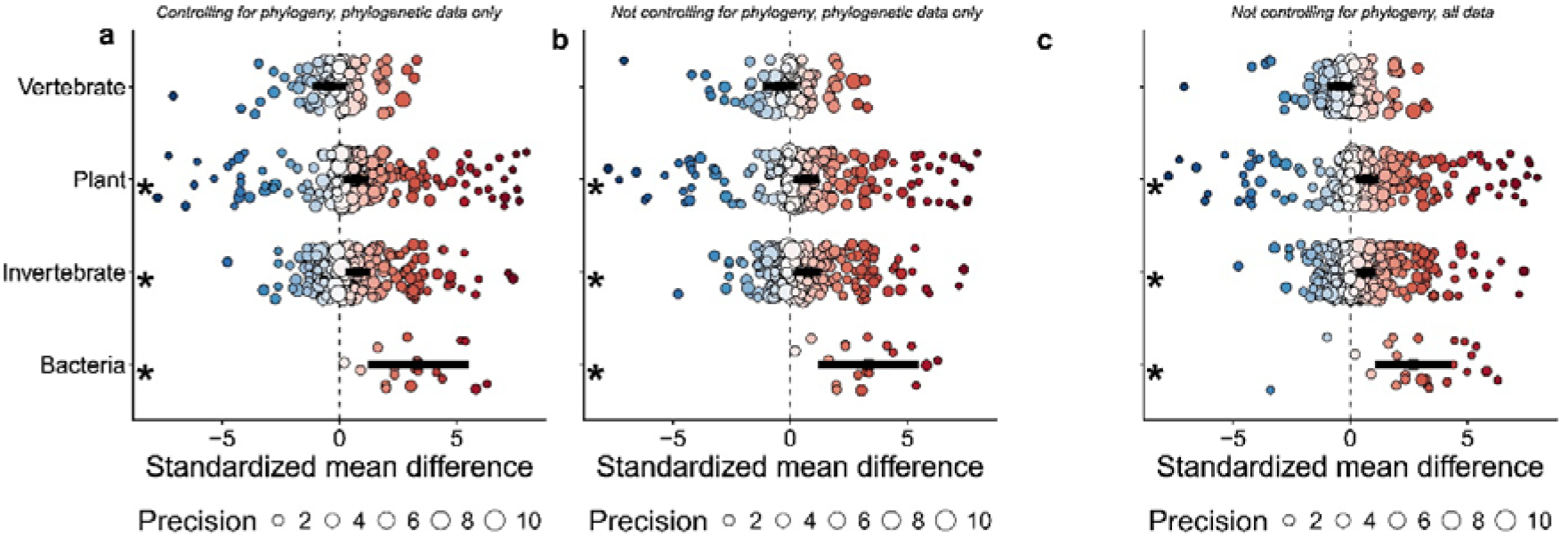
Temperature effects across host taxa. Effects of elevated temperature on parasitism across broad host groups. Columns = model type (phylogenetic, non-phylogenetic, full dataset). Effect sizes are truncated to −8 to 8 (full range: −29.59 to 102.12). Positive values indicate increased parasitism under warming. Black squares show model estimates (±95% CI). Points show individual effect sizes, colored by magnitude and scaled by precision (1/SE). Asterisks denote statistically-significant model estimates.

### Effects of habitat

While the phylogenetically-controlled model estimated a positive (though non-significant) effect (Fig. 4a; Table S3), we found that warming significantly increased parasitism in terrestrial systems in both non-phylogenetic models (all *θ* > 0.6; Fig. 4b-c). No significant temperature-parasite relationships were observed in freshwater or marine systems in any analysis (Fig. 4a-c, Table S3).

**Fig. 4.**
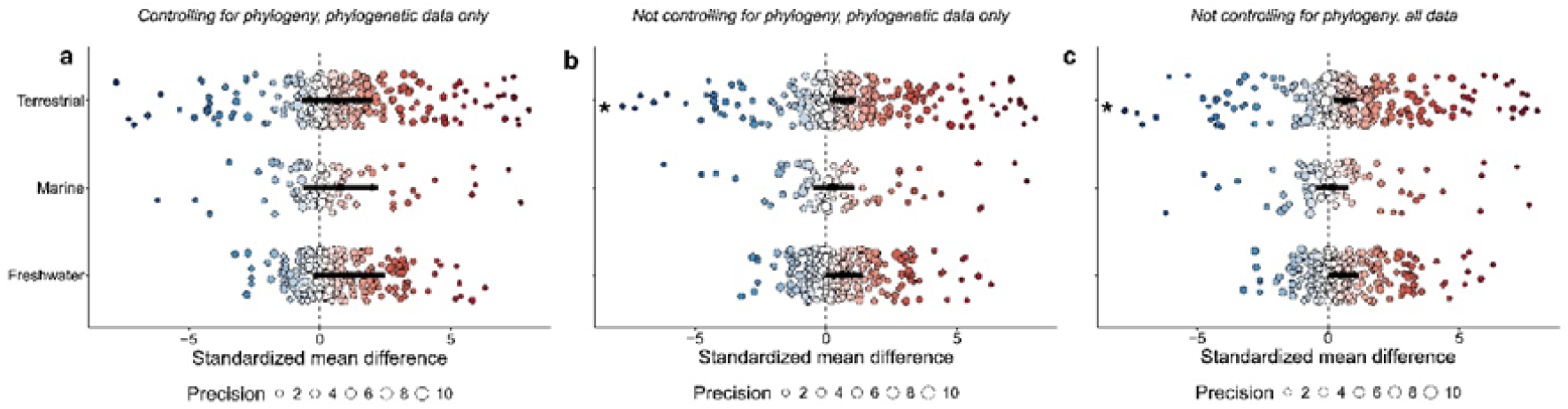
Temperature effects across habitats. Effects of elevated temperature on parasitism in terrestrial, marine, and freshwater systems. Columns = model type (phylogenetic, non-phylogenetic, full dataset). Effect sizes are truncated to −8 and 8 (full range: −29.59 to 102.12). Positive values indicate increased parasitism under warming. Black squares show model estimates (±95% CI). Points show individual effect sizes, colored by magnitude and scaled by precision (1/SE). Asterisks denote statistically-significant model estimates.

### Effects of temperature range

The magnitude of experimental warming (2−30°C) was unrelated to effect size in models excluding viruses (Fig. S1a-b). Including viruses revealed a small negative effect (*θ* = −0.03 [−0.06, −0.004]; Fig. S1c). No significant interactions were detected between temperature range and host type (Fig. S2) or parasite type (Fig. S3), irrespective of phylogenetic correction or viral inclusion.

### Heterogeneity and publication bias

All models exhibited substantial heterogeneity (Table S4), driven primarily by variation among and within studies and by host phylogeny. Funnel plots were asymmetric (Fig. S4), but formal tests revealed no evidence for publication bias (Table S5). Effect sizes were unrelated to sample size (no small-study effect), publication year (no time-lag bias), or statistical significance (no file-drawer effect). Excluding potentially influential outliers did not alter qualitative conclusions in our overall models (phylogenetically-controlled: *θ* = 0.56 [-0.33, 1.45]; non-phylogenetic: *θ* = 0.53 [0.27, 0.79]; including viruses *θ* = 0.49 [0.26, 0.72]). Accordingly, all reported analyses are based on the full dataset, consistent with previous syntheses^8,19^.

## Discussion

Robust tests of the WSWH are crucial for predicting how climate change indirectly affects global health by altering parasites that mediate host health and disease. Our study advances understanding of climate-driven parasitism in several ways. First, previous work has focused on mainly observational studies, providing only phenomenological insights into temperature-parasite relationships. Second, prior syntheses often focused on a limited set of host taxa, reducing the generality of conclusions. By synthesizing exclusively-experimental studies across terrestrial, freshwater, and marine systems, and explicitly accounting for host and parasite phylogeny, our meta-analysis provides the most comprehensive and rigorous test of the WSWH to date.

Across diverse host-parasite systems, elevated temperatures generally increased parasitism. Considering the full dataset, a 1°C rise in temperature corresponded to an average increase in parasitism of 0.19 SD. Non-phylogenetic models detected moderate, statistically-significant increases, while phylogenetically-controlled models yielded directionally-consistent but less precise estimates. These findings indicated that warming frequently favors parasites, yet the responses are evolutionarily-structured and unevenly distributed across the Tree of Life. Importantly, because our synthesis is restricted to controlled temperature manipulations, the observed effects demonstrate that warming alone, independent of shifting species distributions, community composition, or other ecological feedbacks, is sufficient to increase infection in many systems.

### Parasite traits and taxonomic sensitivity

Both macro- and microparasites exhibited positive temperature responses, yet finer taxonomic resolution revealed substantial heterogeneity. Nematodes showed particularly strong and consistent positive responses, likely reflecting their life-history traits. Many nematodes have free-living, environmentally-exposed stages whose development accelerates under warming. For example, several species of the genus *Heterorhabditis* -entomopathogenic nematodes with a free-living environmentally-transmitted larval stage -exhibited elevated infection intensity in the greater wax moth (*Galleria mellonella*) under elevated temperatures^21^. However, strong temperature responses in nematodes are not restricted to environmentally-exposed stages. In *Angiostrongylus vasorum*, infection intensities increased fourfold between 5°C and 15°C due to accelerated parasite growth within gastropod hosts^22^. Elevated temperatures may further increase infection risk by boosting host metabolic and foraging rates, enhancing encounter rates with infective stages^22,23^. These mechanisms – accelerated development, enhanced within-host growth, and increased contact rates - are readily expressed under controlled experimental conditions and likely explain the large effect sizes observed for nematodes. Why we do not see similarly large effect sizes for other parasite groups that also have free-living, environmentally-transmitted stages (such as ticks^24^ or fungi^25^) requires further investigation into the unique life histories of these parasite groups.

Microparasites, which replicate within hosts, also responded positively to warming. For example, oomycete *Phytophthora lateralis* infections on *Rhododendron* leaves produced lesion areas nearly twenty times larger at 28°C than at 2°C^26^. Thus, even parasites confined to host tissues can experience substantial warming-induced increases in infection, although outcomes depend on the balance between parasite performance and host defense.

### Host identity shapes temperature-parasite relationships

Host taxonomy emerged as a key determinant of warming effects. Elevated temperatures consistently increased parasitism in plants, invertebrates, and bacterial hosts, whereas vertebrates exhibited no consistent response and, if anything, weakly negative trends. This contrast likely reflects fundamental differences in immune complexity, physiological buffering, and thermal regulation. Plants and invertebrates rely primarily on innate defenses and lack internal thermal homeostasis, exposing both hosts and parasites to ambient temperature. Similarly, bacteria depend on cell-autonomous resistance and tolerance mechanisms that may be less able to counter accelerated parasite metabolism and replication. Consequently, elevated temperatures can shift the balance in favor of parasites in these systems.

In contrast, vertebrates possess adaptive immune systems capable of mounting temperature-resilient and plastic responses, which may offset temperature-driven increases in parasite growth or transmission. Many vertebrate parasites, particularly endoparasites, develop within relatively stable internal host temperatures, potentially buffering them from external warming. Although behavioral thermoregulation (e.g., habitat selection or microclimate use) is unlikely in laboratory experiments, vertebrate physiological complexity likely constrains warming-induced increases in parasitism. The absence of consistent positive effects in vertebrates underscores the role of physiological regulation and immune plasticity in mediating host-parasite responses to warming.

### Habitat-specific responses

Although positive temperature effects on parasitism occurred across habitats, statistically-significant increases emerged exclusively in terrestrial systems. Because our synthesis is restricted to experimental studies, these patterns cannot be attributed to differences in environmental thermal buffering or field-scale temperature variability between air and water. Instead, they likely reflect biological sensitivity of specific terrestrial host-parasite systems to warming under controlled conditions, rather than intrinsic differences between terrestrial and aquatic environments per se. Moreover, in aquatic systems, thermal manipulations are often constrained by host survival and oxygen availability^27^, which may limit detectable temperature effects.

### Reconciling experimental and observational syntheses

Our findings differ from previous meta-analyses of observational data, which reported weak or inconsistent support for the WSWH^14-16^. Observational field studies integrate direct thermal effects with indirect ecological processes (e.g., host density, behavior, spatial structure), and this ecological realism can obscure direct physiological responses to warming. By focusing exclusively on experimental manipulations, our analysis isolates the direct effects of temperature on parasitism. The contrast between experimental and observational syntheses suggests that direct thermal benefits to parasites may be partially masked in nature by ecological feedbacks. Experiments combining controlled warming with natural ecological complexity will be essential for determining when and where these opposing forces dominate. Such experiments would serve to identify if the direct causal effect of temperature (i.e., the effect of temperature on parasitism *only*, measured in controlled laboratory experiments) differs from the total causal effect (i.e., the effect of temperature on parasitism *plus* other relevant indirect effects manifesting in nature)^28^.

Despite these differences, our results also align with some aspects of previous syntheses. Phylogenetically-controlled models replicate patterns reported by Wolmuth-Gordon et al.^15^, which found no overall positive effect of temperature on parasitism in terrestrial hosts once evolutionary history is accounted for. This attenuation indicates that temperature responses are phylogenetically-conserved, such that closely-related hosts and parasites tend to exhibit similar thermal sensitivities, inflating apparent generality in non-phylogenetic models (like in our analyses). Similarly, Mahon et al.^14^ reported a positive temperature-disease association (which we confirm in the dataset roughly three times larger) when phylogeny is not considered, finding similar effect sizes to what we found. The signal weakens once phylogenetic relationships are incorporated, again suggesting that shared evolutionary history structures host-parasite responses to warming. Together, these independent lines of evidence from multiple syntheses suggest that temperature generally elevates infection, but the strength and detectability of this effect depend on system-specific biology, evolutionary history, and study design.

### Implications and future directions

Despite its breadth, the experimental literature remains geographically and taxonomically biased. Approximately three-quarters of studies derive from temperate systems, with limited representation of tropical taxa (only 5%). Vertebrate and bacterial hosts comprise only ∼18% and 3% of our host taxa, respectively, while fungi, bacteria, protists, and viruses dominate parasite data. Given systematic variation in thermal performance curves and immune strategies across latitude and host clades, expanding experimental work into underrepresented systems is essential for assessing the global generality of the WSWH. Integrating thermal biology with ecological realism, particularly through mesocosms or field-based warming experiments, will be crucial to determine when direct parasite benefits outweigh indirect ecological feedbacks.

### Conclusion

Our synthesis revealed that rising temperatures directly and frequently elevate parasitism across experimentally-studied host-parasite systems. While the magnitude and significance of responses depend on host identity, parasite taxonomy, and habitat, the overarching pattern supports a core prediction of the WSWH: warming tends to benefit parasites more than their hosts. By providing the most comprehensive experimental synthesis to date, our study clarifies a long-standing debate and highlights parasitism as a critical, yet still under-integrated, component of ecological responses to climate change. Because parasites universally affect host fitness, species interactions, and evolution^8,19,29^, incorporating parasitism into predictions of climate change impacts is essential to anticipate the full consequences of a warming world.

## Online Methods

### Literature search and data collection

We conducted a systematic literature search in the Web of Science Core Collection (Basic Search: Topic) up to October 2025 using the following query: *(experiment* OR test* OR expos*) AND temperature AND (parasite* OR pathogen*) AND (prevalen* OR intens* OR infectivity OR transmission OR “parasite fitness” OR infection OR resistan* OR susceptib*) NOT (human OR dairy OR review OR coronavirus OR hospital OR healthcare OR people OR children)*. This search targeted experimental studies quantifying the effects of temperature on parasite prevalence and/or infection intensity. Search terms were defined using an initial screening of 16 benchmark publications representing 10 well-characterized experimental host-parasite systems across 11 journals (1999−2023), ensuring all benchmark studies were retrieved (Supplement 2).

We screened titles and abstracts to exclude studies that did not experimentally-manipulate temperature during host-parasite interactions, were not in English, or were reviews, syntheses, or modelling studies. Full texts were evaluated against inclusion criteria: (i) parasites infected living hosts (including detached plant tissues when these constituted the host), rather than growing in cell cultures, agar plates, water, or dead host material; and (ii) temperature was manipulated during infection, with parasite prevalence and/or intensity measured at a baseline and one or more elevated temperatures. Studies were excluded if parasites were exposed to different temperatures before infection but infectivity was subsequently assessed at a single temperature, hosts were pre-exposed prior to infection (e.g., thermal shock), or infection outcomes were inferred indirectly from host immune, behavioral, physiological, or transcriptomic responses rather than from direct measures of parasite prevalence or intensity. See Fig. S5 for a PRISMA flow chart of our selection process.

When five or more temperatures were tested, we excluded the lowest and highest temperatures, to avoid bias from extreme experimental treatments outside natural thermal ranges. When three to four temperatures were tested, we only extracted data from the highest and lowest temperatures. When parasite metrics were measured at multiple time points using different host individuals (e.g., destructive sampling), each time point was treated as an independent effect size, following previous syntheses^8,19^.

We extracted data on parasite prevalence and/or intensity from text, tables, and supplementary materials, or obtained data directly from authors when necessary. Graphical values were digitized using ImageJ^30^. Standard deviations (SDs) were extracted when reported or were calculated from standard errors (SE) or confidence intervals (CI) using: SD = SE × √N (from SE) and SD = √N × (upper CI − lower CI)/(t value × 2)(from CI)^31^. For box plots, means and SDs were estimated from medians, 25th and 75th quartiles, and extrema using established methods.

### Moderator variables

Parasite prevalence was defined as the proportion of infected hosts per treatment, and infection intensity as the mean parasite burden per infected host. We evaluated multiple biologically-motivated moderators: parasitism metric (prevalence vs. intensity), parasite reproductive strategy (macroparasite vs. microparasite), parasite taxonomic group (virus, trematode, protist, platyhelminth, nematode, fungus, cnidarian, ciliate, bacteria, arthropod), host taxonomic group (vertebrate, plant, invertebrate, bacteria), habitat (terrestrial, freshwater, marine), and temperature difference between control and elevated treatments.

### Effect size

We quantified the effect of temperature on parasitism using Hedge’s *g* (standardized mean difference, SMD), calculated as the difference between control and elevated temperature treatments in SD units. Positive values represent increased parasitism under warming, while negative values indicate reduced parasitism. Hedge’s *g* values of 0–0.3 are considered small, 0.3–0.7 are considered moderate, and > 0.7 are considered large^20^.

### Statistical analysis

We conducted all analyses in R v4.4.3^32^ using the metafor^33^ package. SMDs were calculated for each row of the database using the *escalc* function. Multi-level mixed-effect models were constructed using the *rma*.*mv* function, weighting effect sizes by the inverse of their sampling variance^34^. Categorical moderators (e.g., parasite reproductive strategy, host habitat) were included as fixed effects in separate univariate models.

Because most studies (*n* = 102) contributed multiple effect sizes, we accounted for non-independence by nesting effect size identity within study identity as random effects^35^. We then used a restricted maximum-likelihood estimator to calculate *I*^2^, which estimates the amount of heterogeneity relative to the total amount of variance in the observed effects or outcomes.

### Accounting for phylogenetic relationships

Phylogenetic non-independence among hosts and parasites introduces correlation in comparative analyses^36^, as closely-related taxa may respond similarly to temperature, and host-parasites interactions are shaped by shared (co)evolutionary history^37^. We incorporated host and parasite phylogenies as random effects, including their interaction by calculating the tensor products of the correlation matrices^38,39^.

Because viral parasites and some host-parasite taxa lacked resolved phylogenetic positions, we constructed two datasets and analyzed them using three model structures: (i) phylogenetically-controlled (654 effect sizes from 102 studies), (ii) non-phylogenetic analysis of the same dataset, and (iii) non-phylogenetically-controlled analysis of the full dataset (775 effect sizes from 124 studies). Differences between analyses (i) and (ii) assess phylogenetic structuring; differences between analyses (ii) and (iii) assess the effect of viral parasites and unmapped taxa.

### Publication bias and outliers

We assessed publication bias using multiple approaches^14^: (i) visual inspection of funnel plots (while recognizing their limitations under high heterogeneity^40^); (ii) Egger’s regression to test for small-study effects^40^; (iii) testing for time-lag bias by including publication year as a moderator^41^; and (iv) evaluating the file-drawer problem using the Rosenthal, Orwin, and Rosenberg fail-safe *N* methods, with robustness inferred when fail-safe N exceeded (5 x *N*_*study*_) + 10^40^.

Potentially-influential outliers were identified using Cook’s distance *d*^42^; effect sizes exceeding three times the mean Cook’s distance were removed. We refit intercept-only models to verify if these potential outliers influenced estimates of the temperature effect on parasitism, following prior syntheses^8,19^.

## Data availability

Data and code for this study will be made available after the manuscript has undergone peer review.

## Acknowledgements

We thank Elisabeth Funke for assistance with the literature screening process. This work was funded by a joint Beethoven Life-1 grant from the German Science Foundation (WO 1587/9–1 to JW) and the National Science Centre, Poland (2018/31/F/NZ8/01986 to SC). During the preparation of this manuscript, JW also received funding from the European Union’s HORIZON-MSCA-2022-DN-01 project PHABB (Pathogens of Algae for Biocontrol and Biosecurity; project no. 101120280). AZH benefitted from the musical inspiration of Tony & The Kiki.

## Author contributions

AZH and JW conceived the study. AZH, SC, and JW designed the study. AZH, SC and JW collected data, and AZH performed modelling work and analyzed data. AZH wrote the first draft of the manuscript, and all authors contributed substantially to revisions.

## Notes

### Competing Interest Statement

The authors have declared no competing interest.

