## Supplementary Information for "Warmer world gets sicker – meta-analysis reveals strong increase in parasitism at elevated temperatures across diverse host-parasite systems"

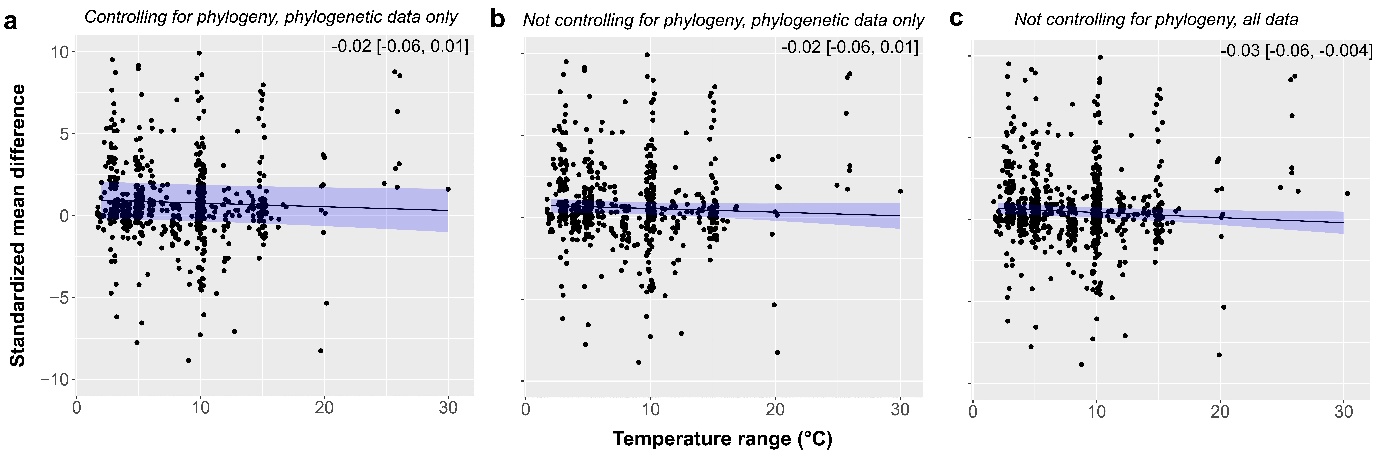
**Fig. S1 – Meta-regression of temperature range.** Relationship between effect size and magnitude of experimental warming (control vs. elevated treatments). Columns = model type (phylogenetic, non-phylogenetic same dataset, non-phylogenetic full dataset). Points show individual effect sizes (horizontally jittered for clarity). Solid lines show model-predicted relationships; shaded ribbons indicate 95% CI. Effect sizes are truncated to -10 to 10. Inset text reports mean effect size and 95% CI.

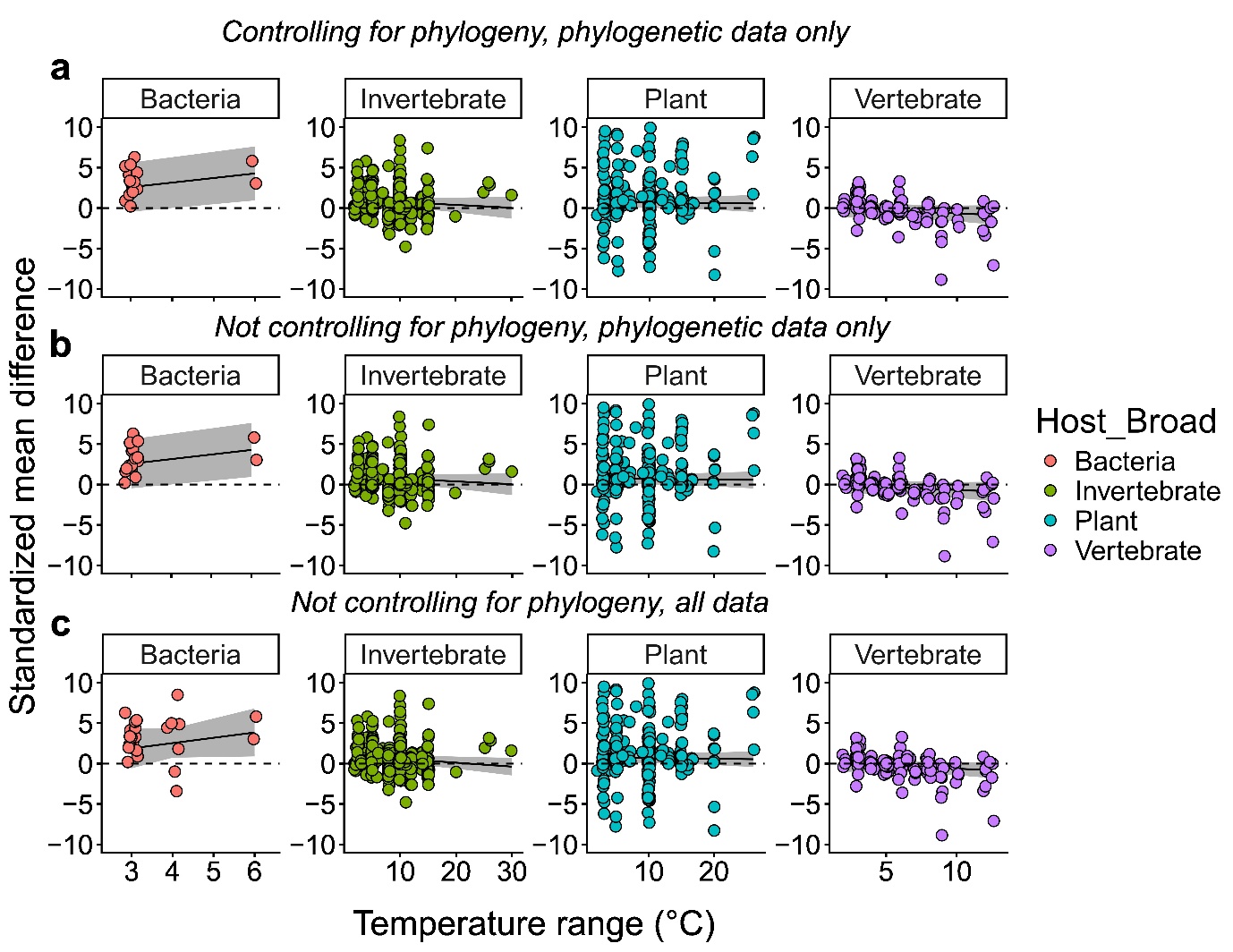

**Fig. S2 – Meta-regression of effect size by host taxa.** Effect size regressed on temperature range (control vs. elevated treatments) across host taxa. Panels = host taxa; rows = model type (phylogenetic, non-phylogenetic same dataset, non-phylogenetic full dataset). Points show individual effect sizes (horizontally-jittered); solid lines show model-predicted relationships; shaded ribbons indicate 95% CI. Effect sizes are truncated to -10 and 10.

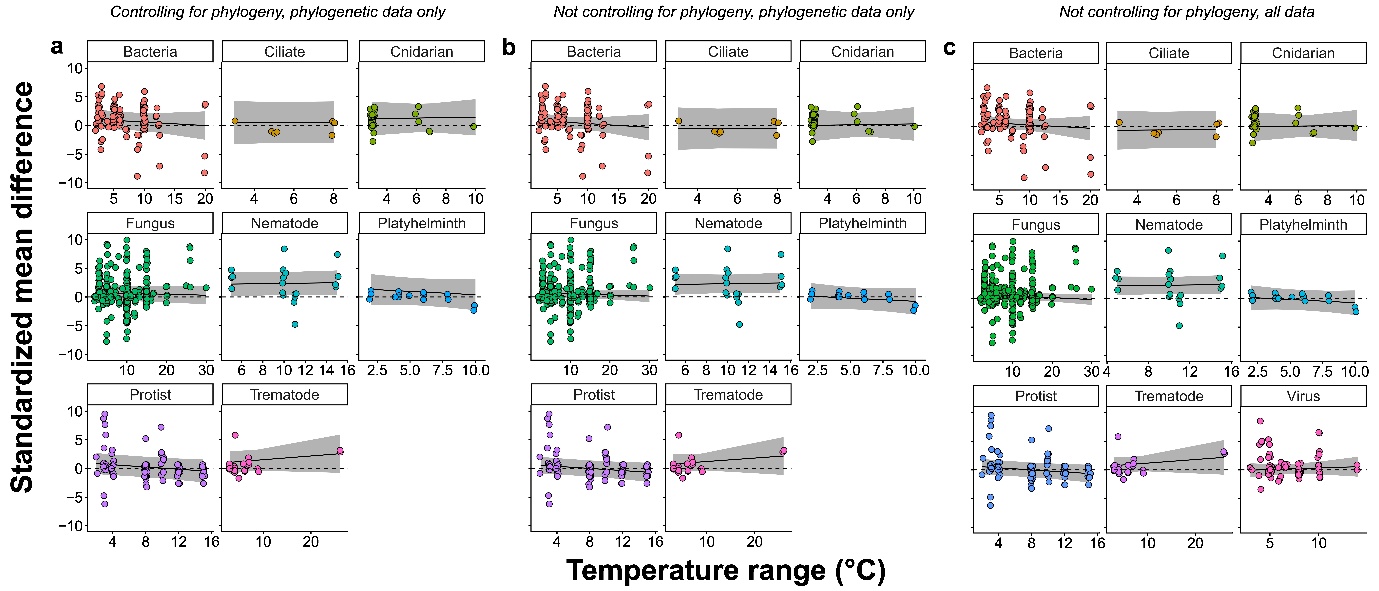

**Fig. S3 – Meta-regression of effect size by parasite taxa.** Effect size regressed on temperature range (control vs. elevated treatments) across parasite taxa. Panels = parasite taxa; columns = model type (phylogenetic, non-phylogenetic same dataset, non-phylogenetic full dataset). Points show individual effect sizes (horizontally-jittered); solid lines show model-predicted relationships; shaded ribbons indicate 95% CI. Effect sizes are truncated to -10 and 10.

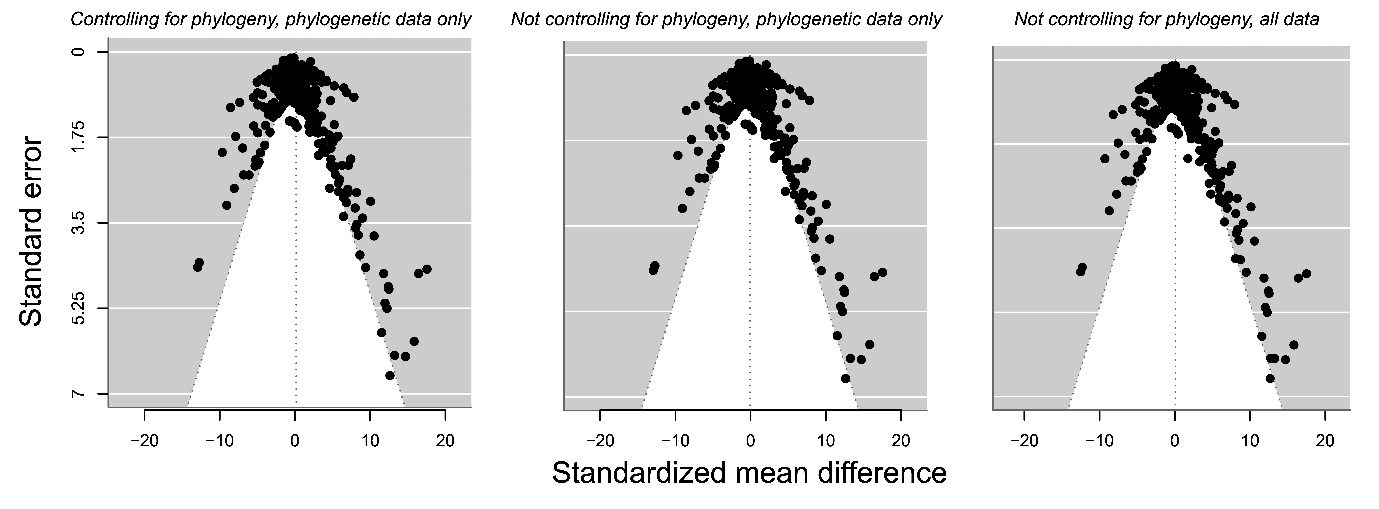

**Fig. S4 – Funnel plots assessing publication bias.** Funnel plots of effect size against standard error. Columns = model type (phylogenetic, non-phylogenetic same dataset, non-phylogenetic full dataset).

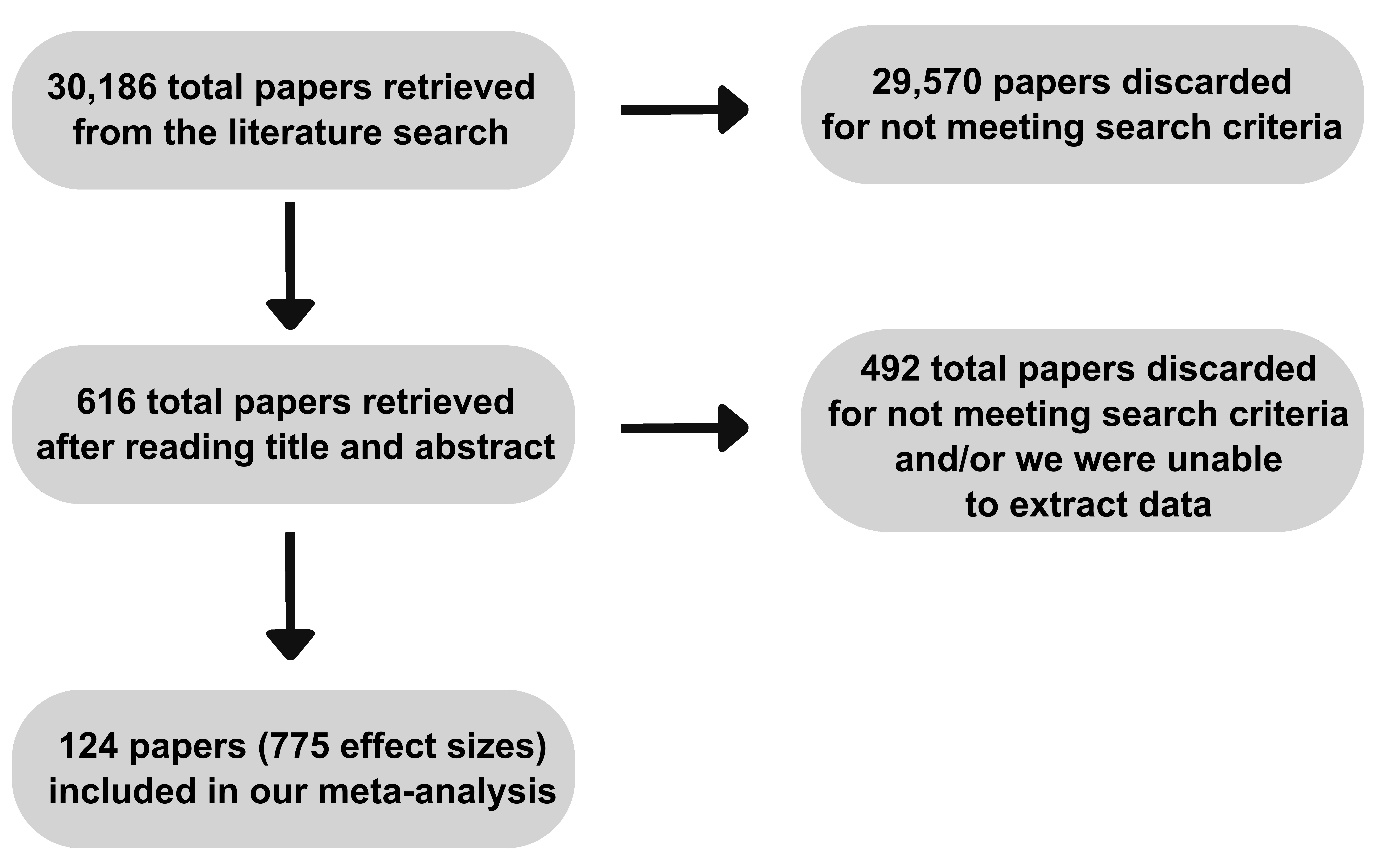
**Fig. S5 – PRISMA flow chart of study selection.** Flow diagram showing the identification, screening, eligibility assessment, and inclusion of studies in the meta-analysis.

**Table S1 – Summary statistics by parasite reproductive strategy and taxon.** Models relating temperature to parasitism across parasite reproductive strategies and parasite taxa. Significant effects are highlighted in **bold**.

| **Parasite reproductive strategy** | | | | |
| --- | --- | --- | --- | --- |
| Dataset and model structure | Factor | Mean effect size | 95% CI | *p* value |
| Controlling for phylogeny, phylogenetic data | Macroparasite | 1.30 | -0.001 – 2.59 | 0.05 |
|  | Microparasite | 0.71 | -0.44 – 1.85 | 0.23 |
| Not controlling for phylogeny, phylogenetic data | Macroparasite | **0.87** | **0.17 – 1.57** | **0.02** |
|  | Microparasite | **0.48** | **0.02 – 0.90** | **0.02** |
| Not controlling for phylogeny, all data | Macroparasite | **0.86** | **0.22 – 1.50** | **0.01** |
|  | Microparasite | **0.41** | **0.07 – 0.74** | **0.02** |
| **Parasite taxonomic group** | | | | |
| Dataset and model structure | Factor | Mean effect size | 95% CI | *p* value |
| Controlling for phylogeny, phylogenetic data | Arthropod | 0.68 | -2.66 – 4.01 | 0.69 |
|  | Bacteria | 0.78 | -0.65 – 2.21 | 0.28 |
|  | Ciliate | 0.56 | -2.83 – 3.95 | 0.75 |
|  | Cnidarian | 1.34 | -1.03 – 3.70 | 0.27 |
|  | Fungus | 0.67 | -0.57 – 1.91 | 0.29 |
|  | Nematode | **2.42** | **0.54 – 4.30** | **0.01** |
|  | Platyhelminth | 0.91 | -1.35 – 3.16 | 0.43 |
|  | Protist | 0.41 | -1.26 – 2.08 | 0.63 |
|  | Trematode | 1.43 | -0.30 – 3.16 | 0.11 |
| Not controlling for phylogeny, phylogenetic data | Arthropod | 0.49 | -2.80 – 3.78 | 0.77 |
|  | Bacteria | 0.49 | -0.43 – 1.40 | 0.30 |
|  | Ciliate | -0.51 | -3.83 – 2.81 | 0.76 |
|  | Cnidarian | 0.13 | -1.88 – 2.13 | 0.90 |
|  | Fungus | 0.54 | -0.02 – 1.10 | 0.06 |
|  | Nematode | **2.33** | **0.86 – 3.81** | **0.002** |
|  | Platyhelminth | -0.30 | -2.12 – 1.51 | 0.75 |
|  | Protist | 0.14 | -1.16 – 1.44 | 0.83 |
|  | Trematode | 1.09 | -0.26 – 2.44 | 0.11 |
| Not controlling for phylogeny, all data | Arthropod | 0.49 | -2.50 – 3.48 | 0.75 |
|  | Bacteria | 0.49 | -0.36 – 1.33 | 0.26 |
|  | Ciliate | -0.52 | -3.54 – 2.50 | 0.74 |
|  | Cnidarian | 0.13 | -1.71 – 1.96 | 0.89 |
|  | Fungus | **0.52** | **0.02 – 1.01** | **0.042** |
|  | Nematode | **2.34** | **0.99 – 3.68** | **0.001** |
|  | Platyhelminth | -0.30 | -1.96 – 1.37 | 0.73 |
|  | Protist | 0.13 | -1.05 – 1.31 | 0.83 |
|  | Trematode | 1.05 | -0.18 – 2.29 | 0.10 |
|  | Virus | 0.17 | -0.62 – 0.95 | 0.68 |

**Table S2 – Summary statistics by host taxon.** Models relating temperature to parasitism across broad host groups.

| Dataset and model structure | Factor | Mean effect size | 95% CI | *p* value |
| --- | --- | --- | --- | --- |
| Controlling for phylogeny, phylogenetic data | Bacteria | **3.35** | **1.20 – 5.49** | **0.002** |
|  | Invertebrate | **0.78** | **0.26 – 1.31** | **0.003** |
|  | Plant | **0.69** | **0.14 – 1.25** | **0.02** |
|  | Vertebrate | -0.44 | -1.18 – 0.31 | 0.25 |
| Not controlling for phylogeny, phylogenetic data | Bacteria | **3.35** | **1.20 – 5.49** | **0.002** |
|  | Invertebrate | **0.78** | **0.26 – 1.31** | **0.003** |
|  | Plant | **0.69** | **0.14 – 1.25** | **0.02** |
|  | Vertebrate | -0.44 | -1.18 – 0.31 | 0.25 |
| Not controlling for phylogeny, all data | Bacteria | **2.71** | **1.08 – 4.33** | **0.001** |
|  | Invertebrate | **0.68** | **0.25 – 1.11** | **0.002** |
|  | Plant | **0.72** | **0.22 – 1.21** | **0.01** |
|  | Vertebrate | -0.42 | -0.99 – 0.15 | 0.15 |

**Table S3 –** Summary statistics by habitats. Models relating temperature to parasitism across terrestrial, freshwater, and marine systems.

| Dataset and model structure | Factor | Mean effect size | 95% CI | *p* value |
| --- | --- | --- | --- | --- |
| Controlling for phylogeny, phylogenetic data | Freshwater | 1.13 | -0.25 – 2.50 | 0.11 |
|  | Marine | 0.79 | -0.65 – 2.22 | 0.28 |
|  | Terrestrial | 0.65 | -0.71 – 2.00 | 0.35 |
| Not controlling for phylogeny, phylogenetic data | Freshwater | 0.68 | -0.05 – 1.41 | 0.07 |
|  | Marine | 0.29 | -0.49 – 1.08 | 0.47 |
|  | **Terrestrial** | **0.65** | **0.15 – 1.16** | **0.01** |
| Not controlling for phylogeny, all data | Freshwater | 0.57 | -0.03 – 1.16 | 0.06 |
|  | Marine | 0.14 | -0.48 – 0.76 | 0.66 |
|  | **Terrestrial** | **0.64** | **0.21 – 1.07** | **0.004** |

**Table S4 – Heterogeneity index (*I^2^*)**. Summary statistics for *I^2^*, quantifying the proportion of total variability in effect size estimates attributable to heterogeneity among true effects.

| Analysis | Heterogeneity index (*I^2^*) |
| --- | --- |
| Controlling for phylogeny, phylogenetic data | I^2^_total_ = 94.27% |
|  | I^2^_between study_ = 33.88% |
|  | I^2^_within study_ = 26.40% |
|  | I^2^_host phylogeny_ = 33.16% |
|  | I^2^_parasite phylogney_ = 2.96 e-06% |
|  | I^2^_host phylogeny x parasite phylogeny_ = 0.83% |
| Not controlling for phylogeny, phylogenetic data | I^2^_total_ = 92.92% |
|  | I^2^_between study_ = 60.24% |
|  | I^2^_within study_ = 32.68% |
| Not controlling for phylogeny, all data | I^2^_total_ = 91.90% |
|  | I^2^_between study_ = 60.74% |
|  | I^2^_within study_ = 31.16% |

**Table S5 – Publication bias and time-lag analyses.** Summary statistics for Egger’s test (meta-regression of effect size on inverse effective sample size [ESS_inverse_]), time-lag bias (meta-regression on publication year), and file-drawer problem (fail-safe *N* test).

| **Publication and time-lag bias** | | | | | | | | | |
| --- | --- | --- | --- | --- | --- | --- | --- | --- | --- |
| Analysis | Variable | | Estimate | | SE | | *Z* | | *p* value |
| Controlling for phylogeny, phylogenetic data | ESS_inverse_ | | -1.46 | | 2.82 | | -0.52 | | 0.61 |
|  | Publication year | | 0.02 | | 0.02 | | 0.84 | | 0.40 |
| Not controlling for phylogeny, phylogenetic data | ESS_inverse_ | | -1.92 | | 2.93 | | -0.66 | | 0.51 |
|  | Publication year | | 0.02 | | 0.03 | | 0.89 | | 0.38 |
| Not controlling for phylogeny, all data | ESS_inverse_ | | -0.73 | | 2.44 | | -0.30 | | 0.77 |
|  | Publication year | | 0.02 | | 0.02 | | 1.03 | | 0.30 |
| **Fail-safe *N*** | | | | | | | | | |
| Method | | Calculated fail-safe *N* | | Threshold value | Pass? |  | |  | |
| Rosenthal | | 147,344 | | 630 | Yes |  | |  | |
| Orwin | | 775 | | 630 | Yes |  | |  | |
| Rosenberg | | 67,246 | | 630 | Yes |  | |  | |
