## Supplement 2 for "Warmer world gets sicker – meta-analysis reveals strong increase in parasitism at elevated temperatures across diverse host-parasite systems"

**Supplement: Validation of search terms.** Document showing 16 benchmark studies known to meet our selection criteria. For each study, the DOI, authors, title, and abstract are listed. Words captured by our search strings are highlighted within the text.

<https://doi.org/10.1002/lno.12257>

Dziuba et al. 2023

Can climate warming save Daphnia from parasites? Reduced parasite prevalence in Daphnia populations from artificially heated lakes

Climate warming might modify infection outcomes and it has been proposed that temperature increase will result in a “sicker world.” We tested this hypothesis by comparing the prevalence of infection in a common freshwater host–parasite system (crustacean Daphnia infected with the ichthyosporean pathogen Caullerya mesnili) between five artificially heated lakes and four nearby non-heated control lakes. The heated lakes, which receive warm water from two power plants, have experienced an elevation in water temperature of ca. 3–4°C for the last 60 yr. Analyses of 5 yr of field data revealed that Daphnia communities from heated lakes had lower parasite prevalence than communities from control sites. To disentangle a possible direct detrimental effect of elevated temperature on the parasite from differences in baseline levels of host resistance, we compared infection susceptibility between Daphnia clones isolated from heated and control lakes, under laboratory conditions at two different temperatures. Daphnia from heated lakes were less susceptible to infection than clones from control lakes, while experimental temperature did not affect infection outcome. The data did not confirm the “warmer hence sicker world” scenario. Instead, it seems that indirect effects of temperature elevation (via shifts in lake hydrology) may restrict its spread into heated lakes. Then, local adaptation to the host from control lakes further inhibits re-establishment of the parasite from control to heated lakes. Our results underline the context-dependency of the impact of temperature increase on host–parasite interactions.

<https://doi.org/10.1111/fwb.13464>

Manzi et al. 2020

Temperature and host diet jointly influence the outcome of infection in a Daphnia-fungal parasite system

Climate change has the potential to shape the future of infectious diseases, both directly and indirectly. In aquatic systems, for example, elevated temperatures can modulate the infectivity of waterborne parasites and affect the immune response of zooplanktonic hosts. Moreover, lake warming causes shifts in the communities of primary producers towards cyanobacterial dominance, thus lowering the quality of zooplankton diet. This may further affect host fitness, resulting in suboptimal resources available for parasite growth.

Previous experimental studies have demonstrated the respective effects of temperature and host diet on infection outcomes, using the zooplankter Daphnia and its microparasites as model systems. Although cyanobacteria blooms and heat waves are concurrent events in nature, few attempts have been made to combine both stressors in experimental settings.

Here, we raised the zooplankter Daphnia (two genotypes) under a full factorial design with varying levels of temperature (the standard 19°C and elevated 23°C), food quality (Scenedesmus obliquus as high-quality green algae, Microcystis aeruginosa and Planktothrix agardhii as low-quality cyanobacteria) and exposed them to the parasitic yeast Metschnikowia bicuspidata. We recorded life history parameters of the host as well as parasite traits related to transmission.

The combination of low-quality cyanobacterial diets and elevated temperature resulted in additive detrimental effects on host fecundity. Low-quality diets reduced parasite output, while temperature effects were context dependent. Overall, we argue that the combined effects of elevated water temperature and poor-quality diets may decrease epidemics of a common fungal parasite under a climate change scenario.

Keywords: climate change, cyanobacteria, disease spread, Metschnikowia, zooplankton

<https://doi.org/10.1098/rsbl.2021.0560>

Schampera et al. 2022

Parasites do not adapt to elevated temperature, as evidenced from experimental evolution of a phytoplankton–fungus system

Global warming is predicted to impact the prevalence and severity of infectious diseases. However, empirical data supporting this statement usually stem from experiments in which parasite fitness and disease outcome are measured directly after temperature increase. This might exclude the possibility of parasite adaptation. To incorporate the adaptive response of parasites into predictions of disease severity in a warmer world, we undertook an experimental evolution assay in which a fungal parasite of phytoplankton was maintained at elevated or control temperatures for six months, corresponding to 100–200 parasite generations. Host cultures were maintained at the respective temperatures and provided as substrate, but were not under parasite pressure. A reciprocal infection experiment conducted after six-month serial passages revealed no evidence of parasite adaptation. In fact, parasite fitness at elevated temperatures was inferior in parasite populations reared at elevated temperatures compared with those maintained under control temperature. However, this effect was reversed after parasites were returned to control temperatures for a few (approx. 10) generations. The absence of parasite adaptation to elevated temperatures suggests that, in phytoplankton–fungus systems, disease outcome under global warming will be largely determined by both host and parasite thermal ecology.

Keywords: adaptation, global warming, host–parasite interaction, disease, cyanobacteria, chytrid

<https://doi.org/10.1017/S0031182018000215>

Agha et al. 2018

Fitness and eco-physiological response of a chytrid fungal parasite infecting planktonic cyanobacteria to thermal and host genotype variation

Understanding how individual parasite traits contribute to overall fitness, and how they are modulated by both external and host environment, is crucial for predicting disease outcome. Fungal (chytrid) parasites of phytoplankton are important yet poorly studied pathogens with the potential to modulate the abundance and composition of phytoplankton communities and to drive their evolution. Here, we studied life-history traits of a chytrid parasite infecting the planktonic, bloom-forming cyanobacterium Planktothrix spp. under host genotype and thermal variation. When expressing parasite fitness in terms of transmission success, disease outcome was largely modulated by temperature alone. Yet, a closer examination of individual parasite traits linked to different infection phases, such as (i) the establishment of the infection (i.e. intensity of infection) and (ii) the exploitation of host resources (i.e. size of reproductive structures and propagules), revealed differential host genotype and temperature × host genotype modulation, respectively. This illustrates how parasite fitness results from the interplay of individual parasite traits that are differentially controlled by host and external environment, and stresses the importance of combining multiple traits to gain insights into underlying infection mechanisms.

Keywords: Host genetic variation; parasite traits; specificity; temperature; transmission; zoospores

<https://doi.org/10.1017/S0031182017002062>

Cuco et al. 2018

Temperature modulates the interaction between fungicide pollution and disease: evidence from a Daphnia-microparasitic yeast model

Temperature is expected to modulate the responses of organisms to stress. Here, we aimed to assess the influence of temperature on the interaction between parasitism and fungicide contamination. Specifically, using the cladoceran Daphnia as a model system, we explored the isolated and interactive effects of parasite challenge (yeast Metschnikowia bicuspidata) and exposure to fungicides (copper sulphate and tebuconazole) at two temperatures (17 and 20 °C), in a fully factorial design. Confirming a previous study, M. bicuspidata infection and copper exposure caused independent effects on Daphnia life history, whereas infection was permanently suppressed with tebuconazole exposure. Here, we show that higher temperature generally increased the virulence of the parasite, with the hosts developing signs of infection earlier, reproducing less and dying at an earlier age. These effects were consistent across copper concentrations, whereas the joint effects of temperature (which enhanced the difference between non-infected and infected hosts) and the anti-parasitic action of tebuconazole resulted in a more pronounced parasite × tebuconazole interaction at the higher temperature. Thus, besides independently influencing parasite and contaminant effects, the temperature can act as a modulator of interactions between pollution and disease.

Keywords: aquatic contamination; copper sulphate; Daphnia spp.; host–parasite interaction; Metschnikowia bicuspidata; multiple stressors; parasitism; tebuconazole

<https://doi.org/10.1098/rsbl.2010.0616>

Temperature effects on parasite prevalence in a natural hybrid complex

Schoebel et al. 2011

Temperature effects on parasite prevalence in a natural hybrid complex Both host susceptibility and parasite infectivity commonly have a genetic basis, and can therefore be shaped by coevolution. However, these traits are often sensitive to environmental variation, resulting in genotype-by-environment interactions. We tested the influence of temperature on host–parasite genetic specificity in the Daphnia longispina hybrid complex, exposed to the protozoan parasite Caullerya mesnili. Infection rates were higher at low temperature. Furthermore, significant differences between host clones, but not between host taxa, and a host genotype-by-temperature interaction were observed.

Keywords: coevolution; Daphnia; genotype-by-environment interaction; host–parasite

<https://doi.org/10.1098/rspb.2005.3404>

Fels and Kaltz 2006

Temperature-dependent transmission and latency of Holospora undulata, a micronucleus-specific parasite of the ciliate Paramecium caudatum

Transmission of parasites to new hosts crucially depends on the timing of production of transmission stages and their capacity to start an infection. These parameters may be influenced by genetic factors, but also by the environment. We tested the effects of temperature and host genotype on infection probability and latency in experimental populations of the ciliate Paramecium caudatum, after exposure to infectious forms of its bacterial parasite Holospora undulata. Temperature had a significant effect on the expression of genetic variation for transmission and maintenance of infection. Overall, low temperature (10 °C) increased levels of (multiple) infection, but arrested parasite development; higher temperatures (23 and 30 °C) accelerated the onset of production of infectious forms, but limited transmission success. Viability of infectious forms declined rapidly at 23 and 30 °C, thereby narrowing the time window for transmission. Thus, environmental conditions can generate trade-offs between transmission relevant parameters and alter levels of multiple infection or parasite-mediated selection, which may affect evolutionary trajectories of parasite life history or virulence.

Keywords: experimental epidemiology, host-parasite interaction, multiple infection, parasite growth

<https://doi.org/10.1046/j.1461-0248.2003.00387.x>

Blandfort et al. 2003

Temperature checks the Red Queen? Resistance and virulence in a fluctuating environment.

Numerous studies have revealed genetic variation in resistance and susceptibility in host– parasite interactions and therefore the potential for frequency-dependent selection (Red Queen dynamics). Few studies, if any, have considered the abiotic environment as a mediating factor in these interactions. Using the pea aphid, Acyrthosiphon pisum, and its fungal pathogen, Erynia neoaphidis, as a model host–parasite system, we demonstrate how temperature can mediate the expression of genotypic variation for susceptibility and virulence. Whilst previous studies have revealed among-clone variation in aphid resistance to this pathogen, we show that resistance rankings derived from assessments at one temperature, are not conserved across differing temperature regimes. We suggest that variation in environmental temperature, through its nonlinear impact on parasite virulence and host defence, may contribute to the general lack of evidence for frequencydependent selection in field systems.

Keywords Coevolution, entomopathogen, evolution of resistance, genetic variation, pea aphid, temperature, virulence.

<https://doi.org/10.1046/j.1365-3032.2003.00309.x>

Stacey et al. 2003

Genotype and temperature influence pea aphid resistance to a fungal entomopathogen

The influence of temperature on life history traits of four Acyrthosiphon pisum clones was investigated, together with their resistance to one genotype of the fungal entomopathogen Erynia neoaphidis. There was no difference among aphid clones in development rate, but they did differ in fecundity. Both development rate and fecundity were influenced by temperature, but all clones showed similar responses to the changes in temperature (i.e. the interaction term was nonsignificant). However, there were significant differences among clones in susceptibility to the pathogen, and this was influenced by temperature. Furthermore, the clones differed in how temperature influenced susceptibility, with susceptibility rankings changing with temperature. Two clones showed changes in susceptibility which mirrored changes in the in vitro vegetative growth rate of E. neoaphidis at different temperatures, whereas two other clones differed considerably from this expected response. Such interactions between genotype and temperature may help maintain heritable variation in aphid susceptibility to fungal pathogen attack and have implications for our understanding of disease dynamics in natural populations. This study also highlights the difficulties of drawing conclusions about the efficacy of a biological control agent when only a restricted range of pest genotypes or environmental conditions are considered.

Key words: Acyrthosiphon pisum, biological control,Erynia neoaphidis, genotypeby environment interaction, resistance, temperature

<https://doi.org/10.1111/j.1420-9101.2008.01555.x>

Vale et al. 2008

Temperature-dependent costs of parasitism and maintenance of polymorphism under genotype-by-environment interactions.

The maintenance of genetic variation for infection-related traits is often attributed to coevolution between hosts and parasites, but it can also be maintained by environmental variation if the relative fitness of different genotypes changes with environmental variation. To gain insight into how infection-related traits are sensitive to environmental variation, we exposed a single host genotype of the freshwater crustacean Daphnia magna to four parasite isolates (which we assume to represent different genotypes) of its naturally co-occurring parasite Pasteuria ramosa at 15, 20 and 25 °C. We found that the cost to the host of becoming infected varied with temperature, but the magnitude of this cost did not depend on the parasite isolate. Temperature influenced parasite fitness traits; we found parasite genotype-by-environment (G × E) interactions for parasite transmission stage production, suggesting the potential for temperature variation to maintain genetic variation in this trait. Finally, we tested for temperature-dependent relationships between host and parasite fitness traits that form a key component of models of virulence evolution, and we found them to be stable across temperatures.

Keywords: cost of parasitism; Daphnia magna; genetic variation; genotype-by-environment interaction; host–parasite; infectivity; Pasteuria ramosa; temperature; transmission; virulence evolution

<https://doi.org/10.1111/j.0014-3820.2005.tb00895.x>

Mitchell et al. 2005

Host-Parasite and Genotype-by-Environment Interactions: Temperature Modifies Potential for Selection by a Sterilizing Pathogen

Parasite-mediated selection is potentially of great importance in modulating genetic diversity. Genetic variation for resistance, the fuel for natural selection, appears to be common in host-parasite interactions, but responses to selection are rarely observed. In the present study, we tested whether environmental variation could mediate infection and determine evolutionary outcomes. Temperature was shown to dramatically alter the potential for parasite-mediated selection in two independent laboratory infection experiments at four temperatures. The bacterial parasite, Pasteuria ramosa, was extremely virulent at 208C and 258C, sterilizing its host, Daphnia magna, so that females often never produced a single brood. However, at 108C and 158C, the host-parasite interaction was much more benign, as nearly all females produced broods before becoming sterile. This association between virulence and temperature alone could stabilize coexistence and lead to the maintenance of diversity, because it would weaken parasite-mediated selection during parts of the season. Additionally, highly significant genotype-by-environment interactions were found, with changes in clone rank order for infection rates at different temperatures. Our results clearly show that the outcome of parasite-mediated selection in this system is strongly context dependent.

Keywords: Daphnia, evolution, Pasteuria, phenotypic plasticity, reaction norm, temperature, virulence

Little et al. 2007

Parasite transgenerational effects on infection

Question: Do past conditions experienced by parasites mediate current levels of infectivity and virulence in the host–parasite combination of Daphnia magna and Pasteuria ramosa? Methods: We varied either temperature (three levels: 15, 20 or 25C) or food supplied to the host (two levels) during a primary infection event, and then harvested parasites and measured their infectivity during a secondary infection event that was subject to the same environmental variation. Result: Past temperatures did not influence any of the infection-related traits measured. By contrast, past food conditions appeared to impact infection, with parasite spores originating from well-fed hosts generally being more harmful. There was no indication that parasites had become specialized to their past environment. Four host genotypes were included in the experiment, and there was evidence that one of four was more sensitive to the environmental history of parasites than were the other hosts, i.e. there was an interaction between host genotype and parasite treatment effects. Conclusion: Overall, parasite transgenerational effects appear to influence the level of harm parasites cause.

Keywords: Daphnia, genetic variation, genotype × environment interaction, maternal effect, parasitism, Pasteuria, pathogen, virulence.

<https://doi.org/10.1111/j.1558-5646.1999.tb05391.x>

Fellowes et al. 1999

Cross-resistance following artificial selection for increased defense against parasitoids in Drosophila melanogaster

An increase in resistance to one natural enemy may result in no correlated change, a positive correlated change, or a negative correlated change in the ability of the host or prey to resist other natural enemies. The type of specificity is important in understanding the evolutionary response to natural enemies and was studied here in a Drosophila-paxasitoid system. Drosophila melanogaster lines selected for increased larval resistance to the endoparasitoid wasps Asobara tabida or Leptopilina boulardi were exposed to attack by A. tabida, L. boulardi and Leptopilina heterotoma at 15°C, 20°C, and 25°C. In general, encapsulation ability increased with temperature, with the exception of the lines selected against L. boulardi, which showed the opposite trend. Lines selected against L. boulardi showed large increases in resistance against all three parasitoid species, and showed similar levels of defense against A. tabida to the lines selected against that parasitoid. In contrast, lines selected against A. tabida showed a large increase in resistance to A. tabida and generally to L. heterotoma, but displayed only a small change in their ability to survive attack by L. boulardi. Such asymmetries in correlated responses to selection for increased resistance to natural enemies may influence host-parasitoid community structure.

Keywords.- Asobaratabida, correlated

responses, Drosophilamelanogaster, encapsulation, Leptopilina, parasitoids, temperature.

<https://doi.org/10.1371/journal.ppat.1000025>

Lazzaro et al. 2008

Genotype-by-environment interactions and adaptation to local temperature affect immunity and fecundity in Drosophila melanogaster.

Natural populations of most organisms harbor substantial genetic variation for resistance to infection. The continued existence of such variation is unexpected under simple evolutionary models that either posit direct and continuous natural selection on the immune system or an evolved life history “balance” between immunity and other fitness traits in a constant environment. However, both local adaptation to heterogeneous environments and genotype-by-environment interactions can maintain genetic variation in a species. In this study, we test Drosophila melanogaster genotypes sampled from tropical Africa, temperate northeastern North America, and semi-tropical southeastern North America for resistance to bacterial infection and fecundity at three different environmental temperatures. Environmental temperature had absolute effects on all traits, but there were also marked genotype-by-environment interactions that may limit the global efficiency of natural selection on both traits. African flies performed more poorly than North American flies in both immunity and fecundity at the lowest temperature, but not at the higher temperatures, suggesting that the African population is maladapted to low temperature. In contrast, there was no evidence for clinal variation driven by thermal adaptation within North America for either trait. Resistance to infection and reproductive success were generally uncorrelated across genotypes, so this study finds no evidence for a fitness tradeoff between immunity and fecundity under the conditions tested. Both local adaptation to geographically heterogeneous environments and genotype-by-environment interactions may explain the persistence of genetic variation for resistance to infection in natural populations.

<https://doi.org/10.1111/j.1461-0248.2007.01146.x>

Laine et al. 2008

Temperature-mediated patterns of local adaptation in a natural plant–pathogen metapopulation

There have been numerous investigations of parasite local adaptation, a phenomenon important from the perspectives of both basic and applied evolutionary ecology. Recent work has demonstrated that temperature has striking effects on parasite performance by mediating trade-offs in parasite life history and through genotype × environment interactions. To test whether parasite local adaptation is mediated by temperature, I measured the performance of sympatric populations against allopatric populations of a fungal pathogen, Podosphaera plantaginis, on its host Plantago lanceolata, across a temperature gradient. I used data on parasite life history and epidemiology to derive fitness estimates to measure local adaptation. The results demonstrate unambiguously that trajectories of host–parasite co-evolution are tightly coupled with parasite adaptation to the abiotic habitat, as the strength, and even direction, of local adaptation varied with temperature. Patterns of local adaptation further depended on how parasite fitness was estimated, highlighting the importance of choosing relevant fitness measures in studies of local adaptation.

Co-evolution, fitness components, genotype·environment interactions, host–parasite interaction, infection, local adaptation

<https://doi.org/10.1111/j.1420-9101.2007.01406.x>

Laine 2007

Pathogen fitness components and genotypes differ in their sensitivity to nutrient and temperature variation in a wild plant pathogen association.

Understanding processes maintaining variation in pathogen life-history stages affecting infectivity and reproduction is a key challenge in evolutionary ecology. Models of host–parasite coevolution are based on the assumption that genetic variation for host–parasite interactions is a significant cause of variation in infection, and that variation in environmental conditions does not overwhelm the genetic basis. However, surprisingly little is known about the stability of genotype–genotype interactions under variable environmental conditions. Here, using a naturally occurring plant–pathogen interaction, I tested whether the two distinct aspects of the infection process – infectivity and transmission potential – vary over realistic nutrient and temperature gradients. I show that the initial pathogen infectivity and host resistance responses are robust over the environmental gradients. However, for compatible responses there were striking differences in how different pathogen life-history stages and host and pathogen genotypes responded to environmental variation. For some pathogen genotypes even slight changes in temperature arrested spore production, rendering the developing infection ineffectual. The response of pathogen genotypes to environmental gradients varied in magnitude and even direction, so that their rankings changed across the abiotic gradients. Hence, the variable environment of spatially structured host–parasite interactions may strongly influence the maintenance of polymorphism in pathogen life-history stages governing transmission, whereas evolutionary trajectories of infectivity may be unaffected by the surrounding environment.

Keywords: coevolution; genotype-by-environment interaction; host–parasite interaction; parasite fitness; Plantago lanceolata
